# Lipid droplet phase transition in freezing cat embryos and oocytes probed by Raman spectroscopy

**DOI:** 10.1101/275164

**Authors:** K.A. Okotrub, V.I. Mokrousova, S.Y. Amstislavsky, N.V. Surovtsev

## Abstract

Embryo and oocyte cryopreservation is a widely used technology for cryopreservation of genetic resources. One challenging limitation of this technology is the cell damage during freezing associated with the intracellular lipid droplets. We exploit a Raman spectroscopy to investigate the freezing of cumulus-oocyte complexes, mature oocytes and early embryos of a domestic cat. All these cells are rich in lipids. The degree of lipid unsaturation, lipid phase transition from liquid-like disordered to solid-like ordered state (main transition) and triglyceride polymorphic state are studied. For all cells examined, the average degree of lipid unsaturation is estimated about 1.3 (with ±20 % deviation) double bonds per acyl chain. The onset of the main lipid phase transition occurs in a temperature range from −10 to +4 °C and does not depend significantly on the cell type. It is found that lipid droplets in cumulus-oocyte complexes undergo an abrupt lipid crystallization, which not completely correlate with the ordering of lipid molecule acyl chains. In the case of mature oocytes and early embryos obtained *in vitro* from cumulus-oocyte complexes, the lipid phase transition is broadened. In frozen state lipid droplets inside the cumulus-oocyte complexes have higher content of triglyceride polymorphic β and β′ phases (∼66%) than it is estimated for the mature oocytes and the early embryos (∼50%). For the first time, to our knowledge, temperature evolution of lipid droplets phase state is examined. Raman spectroscopy is proved as a prospective tool for *in situ* monitoring of lipid phase state in single embryo/oocyte during freezing.

## INTRODUCTION

Cryopreservation of preimplantation embryos and gametes is an important tool, routinely used to back up and exchange laboratory animal strains (1,2) and farm animal breeds (3,4). More recently, some successful examples to apply these technologies and Genome Resource Bank (GRB) concept for endangered animal species were reported (5-7). Despite significant progress in the development of cryopreservation approaches over the last few decades, these approaches can be effectively applied to the relatively small number of mammalian species (8,9). To expand the applicability of GRB concept for mammalian species preservation, further investigations of the factors affecting the cell survival through the procedures of freezing/cryopreservation are needed.

Change of lipid phase state in cells during freezing is among the most important factors limiting the cryopreservation of embryos and oocytes. A number of mammalian species are challenging for embryos/oocytes cryopreservation due to the excessive lipid droplet (LD) content (10-12). The problem of high lipid content in oocytes and embryos seems to concern virtually all the representatives of the *Carnivora* order (5,12), among which a large proportion are endangered species. At least, 28 of 38 extant species of *Felidae* family are either already considered as endangered/vulnerable or will get this status in the nearest future. Embryo cryopreservation was successfully applied to about five felid species (13). However, cryopreservation of oocytes is a big challenge even for the domestic cat (5,14). Modern approaches developed to pass over the lipid caused limitations are based on preliminary remove of LDs from the cell (“delipidation” or “delipation”) (15) or lipid redistribution inside of the cell (so-called “polarization”) (16). Delipidation procedure might be performed mechanically using micromanipulators with a miniature suction pipette (15) or by addition of lipolytic chemicals (17,18). In some cases, these methods help to increase cryotolerance and make possible successful freezing and thawing of the treated samples. For example, pig preimplantation embryo cryopreservation became possible after the introduction of delipidation procedure before freezing (15,19). Domestic cat oocytes were successfully cryopreserved after polarization procedure (16). However, some studies evidence that using delipidation to modulate the lipid content of oocytes, may compromise further embryo development after thawing (16,20).

Development of alternative approaches to increase cryotolerance of the oocytes/embryos with high lipid content needs knowledge about lipid phase states in the freezing samples and about mechanisms of lipid caused cryoinjury. At physiological conditions, lipid state in cells corresponds to liquid-like disordered phase (21). Disordered phase provides a fast diffusion of lipids, lipophilic admixtures in the lateral direction (21,22) and a high permeability of water and water-soluble molecules across phospholipid membranes (23,24), that is essential for cell signalling and transport functions. Membrane proteins are also believed to be sensitive to lipid environment and lipid phase state (21).

Freezing membranes and LDs undergo a lipid phase transition (LPT) into solid-like ordered phases (25,26). Triglycerides contained in the LDs can exist in several polymorphic solid-like states with different phase stability and melting temperatures (26-28). Two different order parameters can be used to characterize the lipid state during freezing. The first one is the degree of acyl chain ordering which can be referred to the number of gauche conformations in the hydrocarbon chain (29). The second order parameter is needed to characterize the crystallization process and related to the translational order in lipid molecules arrangement. In the case of lipid mixtures, these two parameters might change independently, and intermediate phase states, such as liquid ordered state, become possible. In the liquid ordered state, the lipid acyl chains are in an ordered conformational state, but the molecules are arranged randomly, as in a liquid state (30,31). Biological lipid structures consist of multicomponent lipid mixtures. As a result, the description of the phase transition becomes complicated. The simultaneous coexistence of the several phases can be observed (32), and phase transition may occur via intermediate states (33).

Lipid related cryoinjury can arise from the LPT itself or failures in cell regulation caused by ordered lipid state. The LPT is believed to occur at relatively high temperatures and can be responsible for injuries of embryos and oocytes at so-called “chilling” temperatures (about 0 ÷ 10 °C) (34-37). The efficiency of polarization and delipidation procedures drops a hint that cryoinjury arises not from LDs degradation itself but from other cell components somehow related to LDs. Besides energy storing, LDs are known to participate in the regulation of oxidative stress and protein handling (38). At the LPT temperatures, LDs may release of compounds in cell cytoplasm leading to toxic effects or a disturbance in cell regulation mechanisms. Another hypothesis of damaging effect considers the LPT induced phase separation and redistribution of different lipophilic compounds inside of freezing LDs and membranes (39). In the case of triglycerides, the formation of different polymorphic forms can lead to triglyceride separation in LDs.

Nowadays, the mechanisms of cryoinjuries induced by lipids remain obscure, not least because of deficiency of experimental data on lipid state in freezing cells. Electron microscopy observations indicate phase separation in LDs inside of frozen oocytes (40). Arav et al. applied infrared (IR) spectroscopy to investigate temperatures of the LPT in bovine, ovine and human oocytes (34-36). It was found that the LPT for these mammalian species occurs at temperatures above 0 °C and depends on the composition of LDs. Cells with a phase transition at low temperatures are supposed to have a higher tolerance to chilling.

Raman spectroscopy is a prospective approach for contactless *in situ* study of freezing cells with high spatial resolution. In the last decade, this approach was applied to investigate the distribution of ice, cryoprotectant and eutectic crystallisation products in freezing samples (41-44). Moreover, resonance Raman spectroscopy revealed changes of cytochrome redox state in freezing cells (45). The capability of Raman spectroscopy to investigate lipid phase state is proven by studies of frozen cells (2,46) and model lipid systems (47,48).

The present study aimed to identify phase states and transitions occurring in LDs of domestic cat embryos and oocytes during freezing. We investigated stretching CH, C=O, CC Raman bands in the spectra measured from LDs in a wide temperature range to extract the degree of lipid unsaturation, the LPT parameters and polymorphic phase content in a frozen state. A question whether different cell types undergo the LPT in a similar way or not is considered.

## MATERIALS AND METHODS

### Sample preparation

Ovaries and epididymises from domestic cats were obtained after routine ovariohysterectomy and orchiectomy from local veterinary clinics, and were transported to the laboratory within 3–4 h at +4 °C in HEPES buffered TCM-199 (Thermo Fisher Scientific, MA) supplemented with streptomycin (100 μg/ml) and penicillin (100 IU/ml).

The ovaries were minced and cumulus-oocyte complexes (COC) collected into TCM-199 (Thermo Fisher Scientific, MA), supplemented with 5.67 mM HEPES, 25 mM NaHCO_3_, 2.2 mM pyruvate, 2.2 mM sodium lactate, 100 μg/ml streptomycin, 100 IU/ml penicillin and 3 mg/ml bovine serum albumin at 38 °C. Oocytes with uniformly dark ooplasm surrounded by several layers of cumulus cells were rinsed three times in HEPES buffered TCM-199 and cultured in 50 μl of TCM-199 (Thermo Fisher Scientific, MA), containing 5 IU/ml human chorionic gonadotropin (hCG) (Chorulon, Intervet International B.V., the Netherlands), 1 IU/ml equine chorionic gonadotropin (eCG) (Follimag, Mosagrogen, Russia), and supplemented with 2.2 mM sodium lactate, 2.2 mM pyruvate, 25 mM NaHCO_3_, 100 μg/ml streptomycin, 100 IU/ml penicillin and 3 mg/ml BSA under mineral oil at 38 °C, v/v 5 % CO_2_ in 24 hours until the metaphase II stage is reached (*in vitro* maturation, IVM).

For *in vitro* fertilization (IVF), MII oocytes were rinsed three times in Ham’s F-10 (Sigma Aldrich, MO) supplemented with 5 v/v % fetal calf serum, 1 mM L-glutamine, 10 μg/ml heparin, 100 μg/ml streptomycin and 100 IU/ml penicillin and then co-incubated with 10^6^ motile epididimal spermatozoa/ml in 50 μl droplets of IVF-medium under mineral oil in 5 v/v % CO_2_ at 38°C.

Embryos were cultured in Ham’s F-10 (Sigma Aldrich, MO) supplemented with 5 v/v % fetal calf serum, 1 mM L-glutamine, 100 μg/ml streptomycin and 100 IU/ml penicillin at 38 °C, 5 v/v % CO_2_ under oil for up to two days, when the 2-4-cell stage is reached.

To provide Raman study from one to three cells (COCs, mature oocytes or embryos) were transported in plastic straws filled with Ham’s F-10 solution. Before freezing, oocytes/embryos were transferred to cryoprotectant solution of Dulbecco’s Phosphate Buffer Saline (DPBS) and 10 v/v % glycerol. Equilibration with cryoprotectant solution was performed in several steps: on the first step oocytes/embryos were transferred into three times diluted DPBS/glycerol solution for 5 min; then specimen was put into the 10 μl drop of two times diluted DPBS/glycerol solution. Finally, the cells were transported into the undiluted DPBS/glycerol solution and placed on the glass with a cavity. The sample was covered with a piece of mica slice and sealed with paraffin.

### Sample freezing

We carried out experiments with COCs and mature oocytes (three experiments per group) and four experiments with preimplantation embryos (see photos in Fig. 1 a-d). Samples with cells were placed in FTIR600 cryostat (Linkam, UK) cooled by liquid nitrogen vapour flow. Freezing protocol was chosen close to standard slow program freezing protocol conventionally used for mammalian embryos (2,49,50). The sample was cooled to ice nucleation temperature *T*_n_ = –7 °C at cooling rate 1 °C/min. Ice nucleation was induced by touching the sample with copper wire precooled in liquid nitrogen. After ice formation, the sample was kept at *T*_n_ from 10 to 30 min to provide ice recrystallization. The sample was cooled to –40 °C with cooling rate 0.3 °C/min, then at the rate of 1÷2 °C/min to –70 °C and after that with the rate of 5÷10 °C/min to –180 °C. Sample cooling was paused at specified temperatures to acquire Raman spectra. Local temperature near the freezing cell was verified by Raman spectrum of ice (see Fig S1 in Supplementary Material).

**FIGURE 1.**
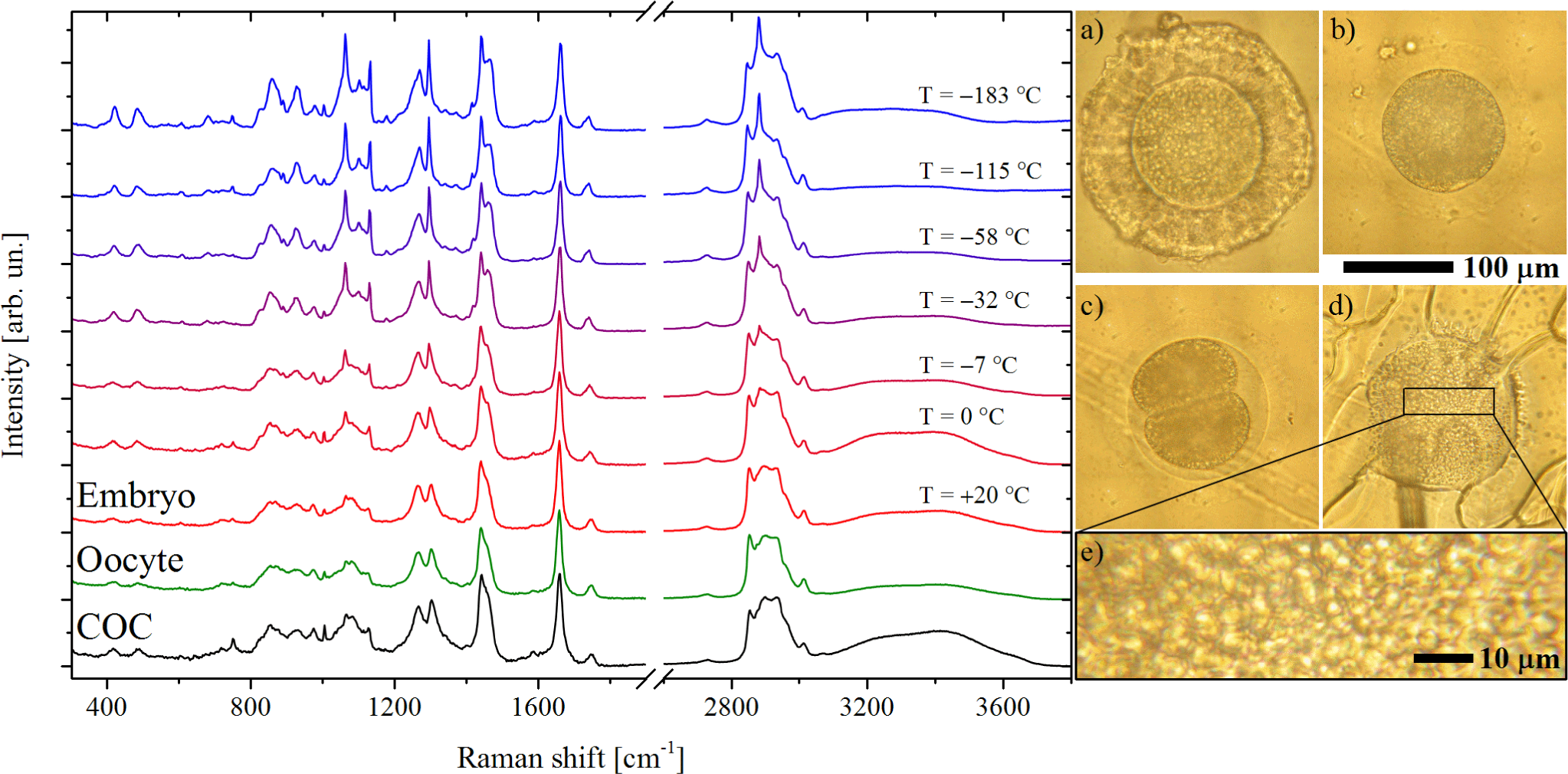
On the left side, representative raw Raman spectra from LDs. Spectra are shifted vertically for illustrative purposes. Order of spectra (bottom to top): COC at +20 °C, mature oocyte at +20 °C, early embryo at +20, 0, –7, –32, –58, –115, –183 °C. Brightfield microscopy photos (on the right side) for a) COC at +20 °C, b) mature oocyte at +20 °C, c,d) embryo at +20, –50 °C, (e) a magnified region with the high amount of LDs.

### Raman experiment

Raman measurements were carried out using a laboratory-built experimental setup (45). Solid-state laser (Millennia II, Spectra Physics) at a wavelength of 532.1 nm was used for Raman scattering excitation. A 100× objective (PL Fluotar L; Leica Microsystems, Germany) with NA=0.75 and working distance 4.6 mm was used to focus laser radiation in approximately 1 μm diameter spot. Irradiation power after objective was 6.5 mW. Scattered radiation was collected using the same objective, and Raman spectra were measured using a monochromator (SP2500i; Princeton Instruments, NJ) equipped with a CCD detector (Spec-10:256E/LN; Princeton Instruments, NJ). Wavelengths for all measured spectra were calibrated using a neon-discharge lamp.

We measured Raman spectra from several substances with known numbers of double bonds. Palmitoleic, linoleic, and linolenic acids, triolein, trillinolein were taken from Sigma Aldrich. Lyophilized phospholipids 1,2-dioleoyl-sn-glycero-3-phosphocholine (DOPC) and 1,2-dilinoleoyl-sn-glycero-3-phosphocholine (DLPC) were taken from Avanti Polar Lipids. Raman spectra were measured from fatty acids and triglycerides in liquid phase state. To measure Raman spectra from phospholipids in liquid-like disordered phase, the suspensions of multilamellar lipid vesicles were prepared using the protocol described previously (48).

The high spatial resolution allows collecting Raman scattering from single LD in the freezing oocyte or the embryo cell (Fig. 1 e). We aimed to measure Raman spectra from the same LD during the experiment. However, it was not always possible due to LD movements and hiding during cell freezing. Thus, we were compelled to change the followed LD one or two times per experiment. Nevertheless, Raman spectra were collected from the neighbouring LDs from the same area inside of the cell. For each experimental point, two spectral ranges were sequentially measured to provide the overall spectral range from 300 to 4000 cm^−1^. For both spectral ranges, several spectra were acquired at each experimental position, followed by spectral averaging. Acquisition time for a single spectrum was 1 min and overall measurement time for one experimental point was 15÷20 min.

## RESULTS

Embryos and oocytes of the domestic cat are rich with lipids, which are mainly found in LDs. Fig. 1 shows representative Raman spectra from LDs in COC, *in vitro* matured oocyte and *in-vitro*-derived early embryo measured at different temperatures. Raman spectra of all cells types contain a similar set of Raman bands. Lipid contribution is manifested by lines of CC stretching vibrations at 1062, ∼1100, 1130 cm^−1^, twisting (1300 cm^−1^) and scissoring (1440 cm^−1^) deformational CH modes, double bonded C=C (1660 cm^−1^) and C=O (1745 cm^−1^) bands. Raman peaks at 2850 and 2882 cm^−1^ manifest the symmetric CH_2_ (sCH) and antisymmetric CH_2_ (aCH) stretching vibrations, respectively. The absence of CN stretching mode at 700 cm^−1^, which is typical for phospholipids (51), indicates that lipid contribution comes mainly from triglycerides and free fatty acids.

In addition to intensive lipid contribution, Raman spectra also contain the lines indicating the presence of proteins and glycerol. The low-intensity line at 1004 cm^−1^ is assigned to phenylalanine contribution, also peaks at 603, 750 and 1586 cm^−1^ correspond to resonance Raman scattering of cytochromes. Other well-known protein lines such as cytochrome peak at 1130 cm^−1^ or Amide I mode at ∼1655 cm^−1^ overlap with lipid lines. The existence of cytochrome Raman lines points on an external contribution from the cytoplasm and the nearest organelles. The protein contribution can be neglected since the intensity of the most intensive protein related peaks (750, 1004, 1586 cm^−1^) in the collected Raman spectra did not exceed 10 % from the deformational band at 1440 cm^−1^ and C=C mode of lipids. Glycerol contribution from LD surroundings has to be taken into account in Raman spectra due to concentration changes resulting from ice formation and cell dehydration. We used glycerol peaks at 420 and 483 cm^−1^ to evaluate the intensity of glycerol peaks (52) and subtract the Raman spectra of aqueous glycerol solution from the raw spectra measured from the cells. Glycerol spectrum is temperature dependent (46) thereby Raman spectra from the cell and glycerol solution measured at same temperatures were applied in subtraction procedure (see details in Supplementary Material).

We used Raman spectra to estimate the degree of lipid unsaturation and to investigate the LPT. To evaluate the degree of unsaturation, Raman intensities of CH deformation mode (CH_2_ scissoring and CH_3_ antisymmetric bending vibrations) and C=C peak were studied. Lipid lines demonstrate pronounced temperature dependence (see Fig. 1). The C=O, CC and CH stretching vibrations are sensitive to the LPT, however, these bands reflect different aspects of lipid structure. The quality of the measured spectra is sufficient for comprehensive analysis of the LPT using all these three spectral regions.

### Analysis of lipid unsaturation degree

The degree of lipid unsaturation is known to affect the temperature of the main LPT. Unsaturation degree can be characterized by the ratio between number of C=C bonds and number of CH_2_+CH_3_ groups (N_C=C_/N_CH2+CH3_). The intensity ratio between C=C peak (*I*_C=C_) and deformational mode at 1440 cm^−1^ (*I*_δCH_) can be used to estimate the degree of lipid unsaturation (53,54). To do so, we constructed a calibration curve (Fig. 2 a) based on several measured triglycerides, phospholipids and free fatty acids with different amount of double bonds per acyl chain (see Fig. 2 b). The N_C=C_/N_CH2+CH3_ was calculated taking into account C=C, CH_2_, CH_3_ groups from acyl chains only. All spectra were measured from samples in disordered phase state at room temperature (+25 °C), which is important because *I*_C=C_/*I*_δCH_ intensity ratio depends on temperature and phase state (for example, see temperature evolution of Raman spectra in Fig. 1). The obtained calibration curve is shown in Fig. 2 b.

**FIGURE 2.**
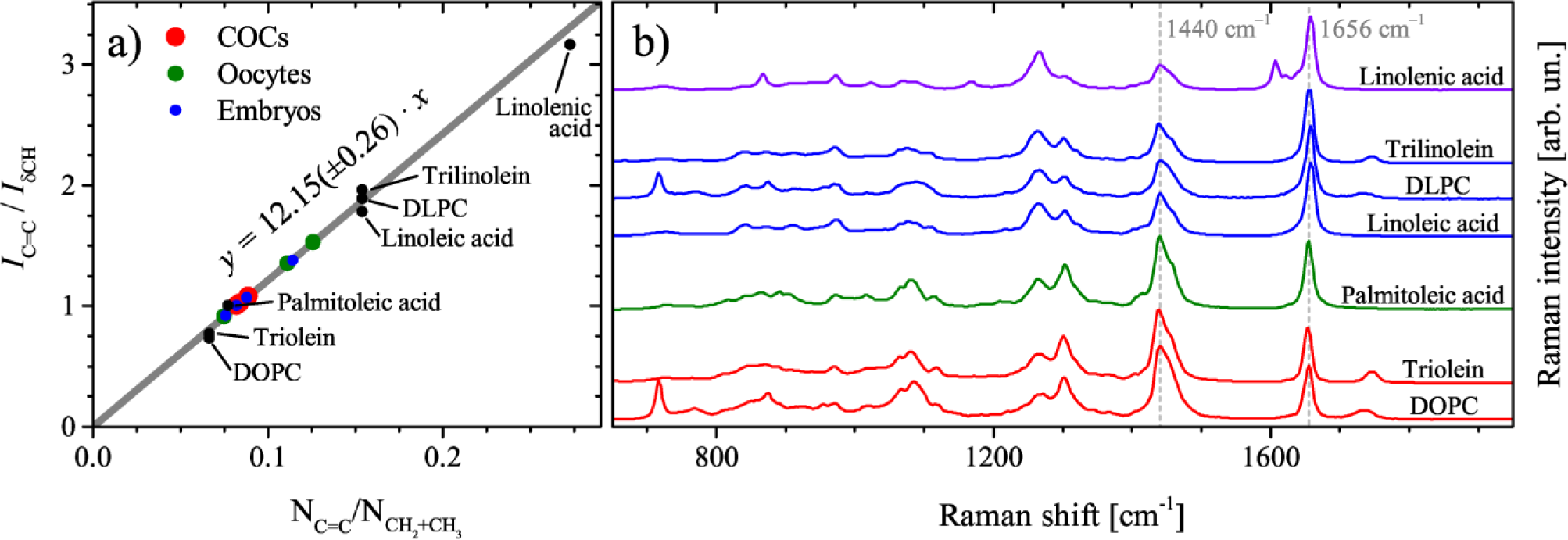
(a) Calibration curve for the evaluation of the degree of unsaturation (N_C=C_/N_CH2+CH3_) from intensity ratio Raman data *I*_C=C_/*I*_δCH_. Gray line is a linear fit (R^2^=0.99). (b) Raman spectra from unsaturated lipids used to calibrate the unsaturation degree in LDs. Spectra are shifted vertically for illustrative purposes.

To estimate the degree of unsaturation, we used spectra of LDs measured at +20 °C. Averaged over all the cells studied *I*_C=C_/*I*_δCH_ is about 1.125 with a standard deviation of 20%. This ratio corresponds to N_C=C_/N_CH2+CH3_=0.0925 or, putting it in other words, ∼1.3 double bonds per typical C18 acyl chain in average. Our experiments do not reveal a significant difference in the degree of lipid unsaturation for different cell types (for details see Table 1). However, the valuable deviation between different experiments were found. The spread in *I*_C=C_/*I*_δCH_ can be partly associated with the systematic experimental errors, i.e. variations in polarization conditions of Raman experiment or protein contribution to measured Raman spectra. Also, N_C=C_/N_CH2+CH3_ spread may come from unspecified parameters such as cat breed or cat diet.

**TABLE 1.** Summary of Raman study results.

| # | $I_{C=C}/I_{\delta_{CH}}$ | $N_{C=C}/N_{CH_2+CH_3}$ | $T^*, ^\circ\text{C}$ | $T_C, ^\circ\text{C}$ | $\beta$ phases fraction |
| --- | --- | --- | --- | --- | --- |
| COC #1 | 1.08 | 0.089 | -4 | -29 ÷ -24 | 65±5% |
| COC #2 | 1.02 | 0.084 | -2 | -7 ÷ -4 |  |
| COC #3 | 1 | 0.082 | +1 | -20 ÷ -13 |  |
| Ooc. #1 | 1.53 | 0.126 | -2 | -30 ÷ -25 | 52±5% |
| Ooc. #2 | 0.91 | 0.075 | -2 | -29 ÷ -8 |  |
| Ooc. #3 | 1.35 | 0.111 | -1 | -62 ÷ -8 |  |
| Emb. #1 | 0.92 | 0.076 | -2 | -33 ÷ -10 | 47±5% |
| Emb. #2 | 1.38 | 0.114 | +4 | -34 ÷ -14 |  |
| Emb. #3 | 1.07 | 0.088 | -10 | -20 ÷ -10 |  |
| Emb. #4 | 1 | 0.082 | -5 | -51 ÷ -30 |  |

### The CH stretching band

Fig. 3 shows the temperature evolution of the CH band which is sensitive to acyl chain conformations and intermolecular interactions. The most striking effect associated with the changes in the lipid molecules state caused by temperature decrease is the intensity increase of aCH mode at 2882 cm^−1^. To investigate the changes in the intensity of aCH we studied intensity ratio between aCH and sCH modes (*I*_aCH_/*I*_sCH_). At high temperatures (*T* > 0 °C), the aCH peak is broadened and *I*_aCH_/*I*_sCH_ ratio is low indicating on inhomogeneous broadening and high variance of conformational states of lipid molecules in disordered phase state. A decrease in temperature leads to a narrowing of the aCH peak, which is associated with freezing of the lipid conformational states and an increase in the ratio *I*_aCH_/*I*_sCH_. The abrupt increase of *I*_aCH_/*I*_sCH_ ratio with temperature decrease can be considered as an evidence of the LPT occurrence.

**FIGURE 3.**
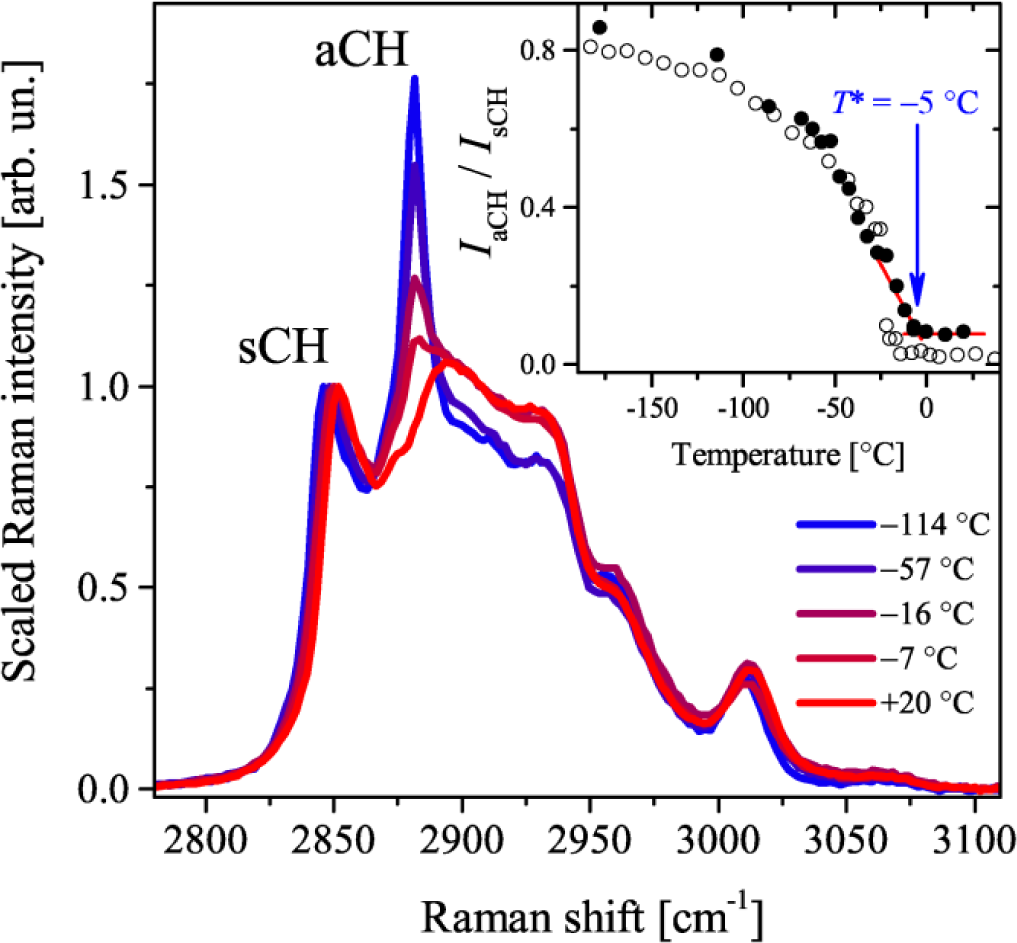
Temperature evolution of Raman CH band from a LD *in vivo* (embryo #4 in Table 1). The inset shows the temperature dependence of *I*_aCH_/*I*_sCH_ ratio for the LD (filled circles) and DOPC vesicles data (empty circles) taken from (48). Arrow marks the temperature corresponding to the onset of the LPT in LDs.

The inset in Fig. 3 shows the *I*_aCH_/*I*_sCH_ ratio temperature dependences of synthetic DOPC vesicles and LD inside the preimplantation cat embryo. DOPC has a one double bond per C18 acyl chain, which is comparable to the degree of lipid unsaturation in LDs in cat oocytes and embryos. Above 0 °C the *I*_aCH_/*I*_sCH_ ratio for both samples can be described with a temperature independent constant. The *I*_aCH_/*I*_sCH_ ratio temperature dependence of synthetic DOPC has a sharp gap corresponding to the LPT at –17 °C. Triolein, which contains three C18 acyl chains with one double bond per each, also exhibits an abrupt change in *I*_aCH_/*I*_sCH_ at the LPT temperature (for example, see temperature dependence for triolein in Fig. S7). However, in the case of LDs in freezing embryos and oocytes, the LPT is broadened, and the increase in *I*_aCH_/*I*_sCH_ occurs more gradually without abrupt changes in *I*_aCH_/*I*_sCH_ ratio. In this case, the *I*_aCH_/*I*_sCH_ ratio deviation from high temperature constant can be considered as the onset of the LPT related to inactivation of conformations state of acyl chains. The onset of the LPT can also be detected by the peculiarity in the temperature behaviour of the symmetric CH_2_ mode (Fig. S4, S5, S6). However, the *I*_aCH_/*I*_sCH_ appears to be a more reliable parameter.

The temperature of the LPT onset (*T*\*) demarcates the completely disordered liquid state and intermediate states with a higher degree of ordering (inset of Fig. 3). For all cells studied the *I*_aCH_/*I*_sCH_ ratio increase begins in temperature range from –10 to +4 °C, with average value of –2 °C. Estimated *T*\* does not correlate with cell type or degree of lipid unsaturation (for details see Table 1). However, maximal spread in *T*\* was found for early embryo stage. At low temperature limit, *I*_aCH_/*I*_sCH_ temperature dependences of LDs are similar to synthetic phospholipid samples (48), following the earlier reported study (46).

### The C=O stretching band

In the Raman spectra, ester carbonyl stretching region can provide the insight into the lipid phase organization (27,28). Triglycerides have three polymorphic forms in solid-like ordered phase: α, β′, and β (55). Raman C=O band can be used to distinguish liquid-like disordered state and three polymorphic forms of the ordered phase (56). The C=O band corresponding to liquid state does not demonstrate any pronounced spectral features and can be described using a Gaussian shape centered at about 1750 cm^−1^. In Raman spectra from polymorphic α form, the C=O band also has Gaussian-like shape, but the band position is shifted to lower frequencies when compared to the spectrum of the liquid state. Other polymorphic forms demonstrate more complex shapes of the C=O band. For example, the β phase of triolein demonstrates in Raman spectra two sharp peaks at 1727 and 1744 cm^−^, spectra of β′ phase of triolein have the peaks at 1730 and 1741 cm^−1^ (56). Raman spectra obtained in our experiments have three broad peaks at about 1727.5, 1734 and 1741 cm^−1^. Full set of these lines does not match to any known lipid polymorphic forms. Taking into account that Raman spectra of different phases depend on particular triglyceride studied, identification of particular β and β′ phases only by Raman spectra seems to be an incorrect task for such complex object as a natural LD. Frozen LD can be formed by a mixture of β and β′ phases of different triglycerides. Therefore, for simplicity, further in the text we will use a term “β phases” implying β, β′ or a mixture of these two phases.

To reveal the lipid crystallization (transition from liquid to solid state, related to ordering in molecules arrangement), we followed the position of the C=O band which was evaluated from C=O band fit with Gaussian. Obtained from different experiments temperature dependences of the C=O band position are shown in Fig 4. It can be seen that the temperature dependences from LDs in COCs have a discontinuity, which the temperature dependences from mature oocytes and early embryos do not have. The detected gap was associated with the transition to solid ordered states, i.e. crystallization of lipids. This gap was used to determine the lipid crystallization temperature (*T*_C_). For three COCs measured *T*_C_ ≈ –5, –17, –27 °C i.e. varies significantly from cell to cell. The temperature dependences of mature oocytes and embryos demonstrate the broadened lipid crystallization occurring in the temperature range from −10 to −50 °C. Temperature ranges of phase transformation for oocytes and embryos are shown in Table 1. In some cases, temperature dependences demonstrate both gradual change and a short gap in C=O band position (Ooc. #1 and Emb. #4 in Fig. 4 and Table 1).

**FIGURE 4.**
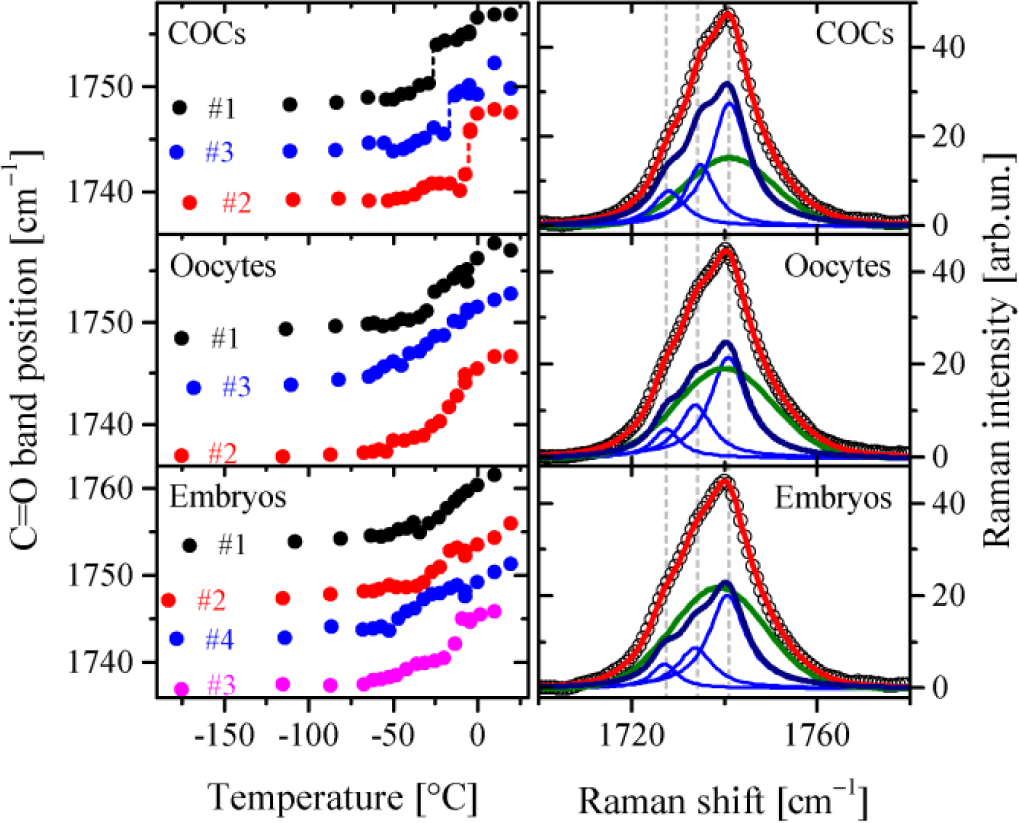
Temperature dependences of C=O band position and band decomposition (at *T* < –60 °C). The left panel shows the temperature dependences of C=O band position for different cell types. Presented temperature dependences are shifted vertically by +5 cm^−1^ for illustrative purposes. Vertical dashed lines in the subpanel with COCs data mark the gaps corresponding to polymorphic transitions. The right panel shows decomposition of experimental C=O spectra on three Lorezians and one Gaussian peak described in the text. Empty circles denote experimental data, the red lines denote the applied fits, the green lines show Gaussian contribution and the navy lines show the contribution from a sum of Lorentz peaks. Vertical dashed lines mark the averaged positions of Lorentz peaks shown by blue lines.

In order to study the phase content of frozen LDs, the spectral shape of the C=O band was investigated. We used spectra with high signal to noise ratio obtained from averaging of the spectra from cells of the same type and all the temperatures below –60 °C. Average spectra were fitted with a sum of three Lorentz peaks and one Gaussian (see Fig. 4). Lorentz peaks simulate contribution from β phases and Gaussian models a contribution from less ordered α phase. When fitting the peak non-negativity constraints were used, the initial values of peaks positions were specified but not fixed. The fit results demonstrated similar parameters of Lorentz peaks (see Fig. S9, S10 and Table S1 in Supplementary Material). Therefore the Lorentz peaks (at 1727.5, 1734, 1741 cm^−1^) do originate from the same structure. The estimated ratio between two phases turned out to be somehow different for COCs and other cells. In LDs of frozen COCs, the portion of β phases is about 65 % (±5 %) from the overall area of the C=O band. Raman spectra of mature oocytes and early embryos reveal about 50 %(±5 %) concertation of β phases. Investigation of C=O band evidences that the freezing of LDs in COCs differs from the freezing of the LDs in mature oocytes and early embryos.

### The C-C stretching region

CC stretching region is widely used in the investigations of acyl chain ordering in model lipid systems (29,48,51,57). Therefore, the capability to investigate CC region in Raman spectra from biological samples was examined. Fig. 5 a shows the temperature dependence of Raman spectra in CC stretching region after baseline and glycerol contributions subtractions. Temperature decrease leads to the intensity growth of the peaks at 1062, ∼1100, 1130 cm^−1^. The mode at the highest frequency at about 1130 cm^−1^, also known as “all-trans” mode, is considered as a reliable measure of all-trans conformations (51). The intensity measurement for CC modes is problematic due to ambiguity in baseline correction and overlap with the cytochrome peak. Therefore, the temperature dependence of the all-trans peak position was examined (see Fig. 5 b). At temperatures above *T*\*, the precision of the parameters evaluation is low due to the low intensity of the all-trans peak. Below this temperature, the all-trans peak becomes sharper and increases the peak position. Further temperature decrease leads to the peak sharpening and the position shift towards to the higher frequencies. Obtained temperature dependence is in qualitative agreement with the data obtained from synthetic lipids such as DOPC (Fig. 5 b). However, low precision of the all-trans peak position estimation at high temperature makes it difficult to detect the LPT using this approach.

**FIGURE 5.**
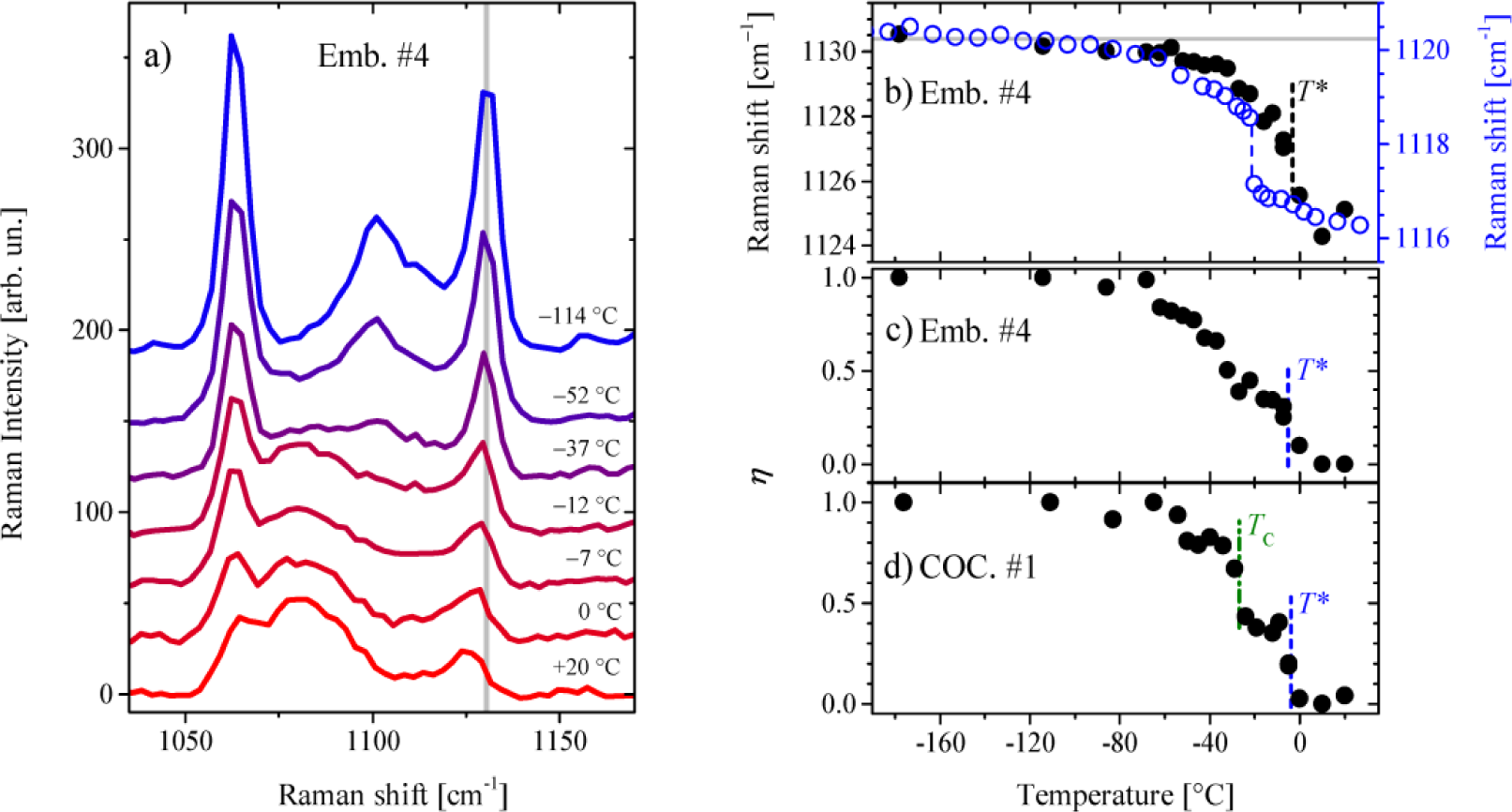
Temperature evolution Raman spectrum in the CC stretching region. Panel a) shows representative stretch CC region of Raman spectra measured from LD *in vivo* (Emb. #4 in Table 1). Gray vertical line denotes the so-called “all-trans” peak. Spectra are shifted vertically for illustrative purposes. On the right, panel b) shows the temperature dependence of the all-trans peak position (Emb. #4). Filled circles represent data obtained from the LD; empty circles correspond to DOPC vesicles data taken from (48). Panels c), d) demonstrate temperature dependences of *η* in the case of Emb. #4 and COC #1, respectively. Vertical dash lines denote *T*\* determined from CH stretching region analyses. Vertical dash-doted line denotes *T*_C_ evaluated from the C=O band analysis.

To study the temperature evolution of the CC stretching region and avoid the problems with all-trans peak analysis, we used a simplified approach based on the CC spectrum linear decomposition into spectral components. This concept is already successfully tested on synthetic lipids (58,59). In approximation that the acyl chains of lipid molecules are in ordered all-trans conformation state at the low-temperature limit (below −100 °C) and the completely disordered at the high-temperature limit (above 10 °C), the CC region was described as the combination of spectral components corresponding to ordered, *S*_*o*_ (*ω*), and disordered, *S*_*d*_ (*ω*), states and linear background. Since this approach involves in the analysis not only all-trans mode, but also other CC modes, more reliable data can be extracted from Raman spectra. Therefore, the Raman spectrum in CC region, *I* (*ω*), was fitted with the linear combination

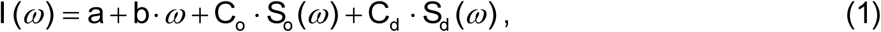

where *a, b* correspond to the parameters of the linear function, *C*_*o*_, *C*_*d*_ are the magnitudes of the ordered and disordered components, respectively. The details of data handling and examples of CC region fits are presented in Supplementary material (Fig. S3). Although the proposed description does not take into account the intermediate conformation states, this approach is sufficient to monitor the most pronounced changes in CC region.

The ratio *η*, defined as *C*_*o*_/(*C*_*o*_ + *C*_*d*_), was used as the acyl chain ordering parameter. The *η =* 0 corresponds to acyl chains in completely disordered conformation state while in the case of the ordered state *η* = 1. The example of *η*(*T*) is shown in Fig. 5 c. The parameter *η* begins to increase at approximately the same temperatures as the *I*_aCH_/*I*_sCH_ ratio. In the case of COC #1, *η*(*T*) demonstrate an abrupt increase at a temperature corresponding to the peculiarity observed in the C=O band (Fig. 4, 5d). However, in the case of COC #3, *η*(*T*) does not show pronounced changes at *T*_C_, which may result from the insufficient quality of measured Raman spectra. For COC #2, *T*\* and *T*_C_ are too close to distinguish these two peculiarities in *η*(*T*) (Fig. S4). Thus, we conclude that the CC region appears to be sensitive to the onset of the LPT related to the ordering of acyl chain conformational states and in some cases it seems to be possible to detect lipid ordering in the molecules arrangement.

## DISCUSSION

Genome Resource Bank (GRB) concept was successfully applied to a number of laboratory and farm animal species (2-4). However, freezing of embryos and oocytes is still challenging for some more exotic mammalian species, especially for those, which oocytes/early embryos are rich with lipids (5,11). Thus to achieve better survival rates and to avoid massive injuries, estimation of the lipid content, evaluation of LPT transition and its control by the conditions of freezing is needed (34-36,46). Here we presented the most accurate and detail data to date about LPT in Felidae oocytes and embryos using contactless Raman approach.

The average unsaturation degree was successfully estimated for the lipids in domestic cat COCs, mature oocytes and early embryos. LDs in domestic cat COCs, oocytes, and early embryos demonstrate a similar degree of unsaturation about 1.3 double bonds per C18 chain and 20 % deviation (N_C=C_/N_CH2+CH3_=0.0925). In comparison, the average unsaturation degree for ovine oocytes, calculated from phospholipid chromatography data (36), is about three times lower (N_C=C_/N_CH2+CH3_=0.0313). The high deviation may result from unspecified parameters since all the oocytes and embryos studied were taken from different cats: breed and diet were not specified. Taking into account the data deviations, we conclude that no drastic changes in the lipid unsaturation degree occur during domestic cat COCs development to mature oocytes and early embryos. At the same time, our results can not exclude the changes in the average unsaturation degree at the level of ∼10 %.

Raman experiment detected the LPT in freezing COCs, mature oocytes and early embryos of a domestic cat. The study of temperature evolution of measured Raman spectra revealed two peculiarities in lipid state in freezing cells. The first one (*T*\*) was observed in temperature dependences of Raman spectra in CH_2_, CC stretch regions. Above *T\** = −2 (−10 ÷ +4) °C, the LD is completely in the liquid disordered state, while below *T*\* the CC and CH bands evidence on the partial ordering of acyl chains. In earlier studies of LPT in oocytes (34-37) and embryos (46), only CH_2_ stretching modes were investigated. Averaged *T*\* is in agreement with the recently reported Raman study of the LPT in mouse embryos, where the LPT was detected in the temperature range from −7 to 0 °C (46). However, for the other species, the LPT is reported at temperatures above 0 °C (34-37). Bovine oocytes undergo the LPT in the temperature range from +13 to +20 °C (35), and for ovine oocytes the broadened LPT occurs at +16 °C (36). Following the conception that chilling injury depends on the LPT (34,36,60), the comparison of different species indicates that embryos and oocytes of domestic cat should have higher chilling tolerance as compared to the ovine or bovine ones. Possibly, this might be the reason that the embryos of the domestic cat were successfully cryopreserved in 1988 (61), much earlier than the embryos of any other Carnivora species. Since then, *in vivo* and *in vitro* produced embryos of the domestic cat were regularly frozen mostly by conventional freezing methods (13,62,63), although very few successful embryo cryopreservation reports were published for other Carnivora animal species (5).

In our interpretation, below *T\** the lipids start to turn from the liquid disordered to the intermediate liquid ordered state. In some cases, mainly in oocytes and early embryos, further cooling is followed by the simultaneous gradual increase in acyl chains ordering and translational ordering in the molecular arrangement. In the case of COCs demonstrating additional peculiarity in the behaviour of C=O mode, rapid ordering in lipid molecules arrangement (i.e. crystallization) occurs at *T*_C_ ≈ −19 (−27 ÷ −5.5) °C. Between *T\** and *T*_C_, LD seems to occur in the intermediate liquid-ordered state. Based on our data on Raman spectroscopy (Fig. S8), we hypothesize that lipids in this intermediate state are ordered in acyl chain conformational state and disordered in the translational arrangement of the molecules. It should be noted that in a homogeneous triglyceride system the ordering of hydrocarbon chains and molecular arrangement occur abruptly at the same temperature (see triolein example in Fig. S7).

Currently, three models describing the liquid-crystalline phase of triglycerides in the disordered state are proposed (27): smectic (64), nematic (65) and discotic (66,67). The last model assumes that in liquid state triglyceride molecules have splayed orientation of acyl chains, forming a discotic (Y-like) conformation state with disordered hydrocarbon chains. The first two models use the concept that lipid molecules in liquid state have a more specific orientation of acyl chains resembling a tuning-fork, which is also known as h-like conformation. Since h-like triglyceride conformation state appears in crystalline phase state, this conformation can be considered as more predisposed to hydrocarbon chains ordering. The smectic phase is supposed to be the most ordered of the three phases mentioned, in this phase the molecules form distinct lamellar structures with a liquid-like translational disorder inside the layers. Therefore, it can be proposed that below *T*,* triglycerides undergo one of the hypothetical transitions: from discotic to nematic, from nematic to smectic (33) phases or even from discotic to smectic phase state. These transitions convert triglycerides into conformational states more suitable for the ordering of acyl chains. The translational ordering of triglycerides (i.e. crystallization) is depressed, since LD consists of multicomponent triglyceride mixture enriched with different admixtures, such as cholesterol. Only at *T*_C_ the phase separation and crystallization of the supercooled mixture takes place.

The detected change in the C=O band position corresponds to triglyceride crystallization. Investigation of the C=O band position indicates on the formation of a mixture of different polymorphic forms. In a frozen state, about 65 % of triglycerides in COCs turn in highly ordered β phases versus 50 % for LDs of matured oocytes and embryos. COCs were taken directly from ovarian tissue, while mature oocytes and early embryos were obtained after *in vitro* procedures (IVM/IVF/IVC). Thus, we suggest that incubation might be the source of the differences between COCs and later stages. Mass-spectrometry study evidence that lipid content may differ in fresh and *in vitro* cultured oocytes and embryos (68,69). These observations are also in agreement with the different biological properties of *in vitro* and *in vivo* matured oocytes (70). Interesting to note that COCs and later stages of development demonstrate the similar degree of lipid unsaturation and *T\**, but different *T*_C_ and crystallized states. Probably, this effect is associated with the changes in the composition of lipophilic admixtures such as cholesterol.

While Raman spectroscopy was already introduced for the investigation of the LPT in early embryos (46), the present study expands the use of this approach to reveal the details of the LPTs in single oocytes and embryos. Raman spectroscopy can be considered as a method of choice for the *in situ* LPT research comprising individual cell monitoring. The capability of individual cell monitoring is especially critical for rare and endangered species. Single cell investigation also can help to avoid the effect of the LPT blurring that inevitably happens in the case of simultaneous study of multiple cells, this is important in the case of sharp transitions (for example see COC data in Fig. 4). Raman experiment can be performed with a high spatial resolution corresponding to the resolution of a confocal microscope and does not suffer from water absorbance limitations. It is noteworthy that the same series of Raman spectra contain information about the degree of lipid unsaturation, the onset of the LPT and triglyceride crystallization in freezing cells. In perspective, contactless label-free Raman approach can be embedded into actual cryopreservation systems and protocols to monitor the phase state of freezing and frozen cells.

## CONCLUSION

In this study, we investigated the phase transitions in the LDs within the frozen COCs, mature oocytes and early embryos of domestic cats using Raman spectroscopy. The specific results of the study are:

i. The average degree of lipid unsaturation (N_C=C_/N_CH2+CH3_) was estimated to be about 0.0925 (with 20% deviations). No significant differences in lipid unsaturation are found between COCs, matured oocytes and preimplantation embryos.
ii. Investigation of the temperature dependences of CC and CH_2_ Raman scattering lines made it possible to detect the onset of the lipid phase transition, which occurs typically at –2 ° C. No significant differences are found between different sample types: COCs, mature oocytes and early embryos. Temperature behaviour of CH_2_ modes in Raman spectra from LDs in all these samples appears to be close to known temperature dependences of CH_2_ stretching modes in synthetic lipid systems. Above the phase transition onset temperature, lipids are in a liquid disordered state in which the LDs can participate in cellular metabolism.
iii. The C=O band is used to reveal triglyceride crystallization in LDs of the cat’s cells during freezing. It is shown that the crystallization of LDs occurs differently for different cell types. COCs undergo a sharp transition, which occurs within the temperature range from –27 to –5 °C. In the case of early embryos and mature oocytes, lipid crystallization occurs gradually during freezing. In our experiments, the composition of polymorphic forms of triglycerides in frozen LDs differs for complexes of COCs and other stages of development.

Finally, we demonstrated that single cell Raman spectroscopy can provide *in situ* label-free characterization of lipid phase transitions in freezing oocytes and embryos. The proposed approach opens the prospects for monitoring of the lipid phase transitions in various lipid-rich embryos and oocytes.

## SUPPORTING MATERIAL

Supporting Material including ten figures and one table is available at http://…

## AUTHOR CONTRIBUTIONS

N.V.S., S.Y.A. and K.A.O designed research; K.A.O and V.I.M. performed the experiments; K.A.O. processed raw data; K.A.O. and N.V.S. analyzed data; K.A.O. wrote the manuscript with contributions from all coauthors.

## ACKNOWLEDGMENTS

We thank V.V. Kozhevnikova for her participation in providing the domestic cat ovaries used in this study and helping.

This work was supported by RFBR (Grant No. 16-04-01221). Part of the experiments was performed in the Multiple-access center “High resolution spectroscopy of gases and condensed matters” in IA&E SBRAS (Novosibirsk, Russia). Biological part of the experiments was performed in the Federal Research Center “Institute of Cytology and Genetics” SB RAS (Novosibirsk, Russia).

